# Light-dependent activation of *TCP4* by the GATA25 transcription factor promotes petiole and hypocotyl elongation in *Arabidopsis thaliana*

**DOI:** 10.64898/2026.10.19.760297

**Authors:** Kihwan Kim, Junseop Shin, Won-Chan Kim

## Abstract

Plants coordinate cell elongation with the daily light/dark cycle, an important adaptive strategy for optimizing growth under changing environmental conditions. However, the transcriptional mechanisms linking light signals to plant developmental growth remain poorly understood. In this study, we identified the GATA25 transcription factor as a regulator of petiole and hypocotyl elongation in response to light and photoperiod. Overexpression of *GATA25* in *Arabidopsis thaliana* increased petiole and hypocotyl lengths and elevated *TCP4* transcript levels. We demonstrated that GATA25 directly binds to the promoter of *TCP4* and activates its transcription using transcriptional activation analysis, a glucocorticoid receptor-based inducible system, chromatin immunoprecipitation, and electrophoretic mobility shift assays. Furthermore, yeast two-hybrid assay and bimolecular fluorescence complementation analyses showed that GATA25 physically interacts with the E3 ubiquitin ligase COP1. Notably, overexpression of GATA25 did not increase *TCP4* transcript levels in dark conditions, suggesting that GATA25 activity is suppressed in dark conditions. Based on these findings, we propose a model in which GATA25 is destabilized by COP1 in dark conditions, restricting *TCP4* activation to the light conditions. Taken together, these findings provide a framework for understanding how the daily light/dark cycle regulates cell elongation in a photoperiod-dependent manner through the GATA25-TCP4 module.

## Introduction

Plants adjust the growth of their cells to daily and seasonal changes in the light environment (Kami et al., 2010). Hypocotyl and petiole elongation are sensitive to light quality, light intensity, and photoperiod, reflecting the coordination of photoreceptor signaling with the circadian clock (Nozue et al., 2007; Niwa et al., 2009). In *Arabidopsis thaliana*, hypocotyl elongation is greater under short-day (SD) conditions than long-day (LD) conditions. This photoperiodic response is explained by an external coincidence mechanism, in which the clock-regulated expression of *PHYTOCHROME-INTERACTING FACTOR 4* (*PIF4*) and *PIF5* coincides with the dark period, when the light-activated photoreceptors no longer repress PIF protein accumulation (Nozue et al., 2007; Niwa et al., 2009). These studies established that the temporal regulation of gene expression and light-dependent protein stability jointly determine growth-promoting activity. However, transcriptional regulators other than PIFs that link the light/dark cycle to cell elongation remain poorly understood.

GATA transcription factors regulate diverse developmental and environmental responses (Luo et al., 2010; Chiang et al., 2012; Schwechheimer et al., 2022; Schröder et al., 2023; Zappone et al., 2026). Among them, GATA25 transcription factor belongs to the ZIM subfamily of GATA transcription factors, which contain a TIFY domain, a CCT domain, and a GATA-type zinc finger domain (Reyes et al., 2004; Shikata et al., 2004). Overexpression of *ZIM,* also known as *GATA25*, promotes hypocotyl and petiole elongation under various light wavelengths (Shikata et al., 2004). In addition, *GATA25* expression is regulated by the circadian clock, with peak transcript abundance around subjective dawn (Manfield et al., 2007), and GATA25 transcription factor has been shown to promote hypocotyl elongation by directly activating *PIF4* transcription (Kim et al., 2023b). These findings suggest that GATA25 transcription factor may link circadian regulation and cell elongation. However, although GATA25 transcription factor has been shown to regulate *PIF4* transcription, whether it also modulates cell elongation through other downstream transcription factors remains unclear. Identifying such targets would clarify how GATA25 transcription factor connects light signals to the developmental programs that drive cell growth.

The TEOSINTE BRANCHED 1/CYCLOIDEA/PROLIFERATING CELL NUCLEAR ANTIGEN FACTOR 4 (TCP4) transcription factor is a plausible candidate for this connection because it promotes hypocotyl elongation and exhibits daily variation in transcript abundance (Challa et al., 2016; Kubota et al., 2017). TCP4 transcription factor belongs to the class II CINCINNATA-like TCP family and is regulated post-transcriptionally by microRNA319 (miR319) (Palatnik et al., 2003; Sun et al., 2017). In addition to promoting the transition from cell proliferation to differentiation during leaf development (Schommer et al., 2014), TCP4 transcription factor directly activates the auxin biosynthetic gene *YUCCA5* (*YUC5*) to promote hypocotyl elongation (Challa et al., 2016). Notably, TCP4 induction did not enhance hypocotyl elongation in dark-grown seedlings, suggesting that its growth-promoting effect depends on the light environment (Challa et al., 2016). Furthermore, TCP4 interacts with PIF4, and these transcription factors cooperatively activate the expression of the cell cycle inhibitor *KIP-RELATED PROTEIN 1* (*KRP1*) to restrict leaf cell proliferation under elevated temperature (Saini et al., 2022). These findings highlight the role of TCP4 in integrating developmental and environmental signals to regulate plant growth. Interestingly, *TCP4* exhibits a daily expression pattern, with transcript levels peaking between Zeitgeber time (ZT) 10 and ZT13 under LD conditions (Kubota et al., 2017). Although transcriptional regulators of *TCP4* have been identified (Li et al., 2016), the upstream mechanisms linking *TCP4* expression to the daily light/dark cycle remain poorly understood. Given the involvement of GATA25 transcription factor in light-responsive growth and its circadian-regulated expression, an important question is whether GATA25 transcription factor directly regulates *TCP4* transcription and whether this regulation is dependent on light conditions.

The potential light dependence of GATA25-mediated *TCP4* regulation raises the question of how photoreceptor signaling might control GATA25 activity at the protein level. Phytochrome B (phyB), a red/far-red light photoreceptor, contributes to the regulation of hypocotyl growth under changing environments (Sellaro et al., 2019; Kim et al., 2020; Song et al., 2021; Zhang et al., 2025). One mechanism linking phyB signaling to transcription factor activity involves the E3 ubiquitin ligase CONSTITUTIVE PHOTOMORPHOGENIC 1 (COP1). COP1 targets positive regulators of photomorphogenesis, including ELONGATED HYPOCOTYL 5 (HY5), for proteasomal degradation (Osterlund et al., 2000; Lau and Deng, 2012). Its nuclear abundance increases in darkness and decreases in response to light (von Arnim and Deng, 1994; Pacín et al., 2014). Light-activated phyB interacts with SUPPRESSOR OF PHYTOCHROME A-105 (SPA) proteins and disrupts their association with COP1, reorganizing and inhibiting the COP1/SPA complex (Sheerin et al., 2015). The inhibition allows COP1 substrates to accumulate and modulate downstream transcriptional responses. The temporal consequences of COP1-mediated turnover are illustrated by CONSTANS (CO), in which COP1 promotes CO degradation at night, contributing to the temporal pattern of CO accumulation underlying photoperiodic flowering (Jang et al., 2008; Liu et al., 2008; Sarid-Krebs et al., 2015). Notably, the C-terminal region of CO containing the CCT domain is required for its interaction with COP1 (Jang et al., 2008; Liu et al., 2008). Because GATA25 also contains a CCT domain (Reyes et al., 2004; Shikata et al., 2004), this shared structural feature provides a rationale for investigating whether COP1 interacts with GATA25 and modulates its ability to activate *TCP4* in a light-dependent manner. Here, we investigated the transcriptional target of GATA25 and the potential regulation of its activity by COP1 to understand cell elongation in response to light and photoperiod. We identified *TCP4* as a direct transcriptional target of GATA25 and compared GATA25-dependent changes in TCP4 expression and hypocotyl elongation under LD and SD conditions. Furthermore, we demonstrated a physical interaction between GATA25 and COP1. These findings identify a GATA25-TCP4 transcriptional module and support a model in which COP1 may contribute to the light-dependent regulation of GATA25 activity.

## Materials and methods

### Plant materials and growth conditions

*Arabidopsis thaliana* (*A. thaliana*) ecotype Columbia-0 (Col-0) was used as the wild type (WT) and as the genetic background of all transgenic plants and mutants in this study. The *GATA25*-overexpressing lines (*GATA25*OX #3 and #13) and the clustered regularly interspaced short palindromic repeat (CRISPR)/CRISPR-associated protein 9 (Cas9)-mediated *gata25* mutant were described (Supplementary Fig. S1-S3). Briefly, the *gata25* mutant was generated with a guide RNA targeting the first exon of *GATA25* (5’- GCCTCCGATTTGATTCCCGA<u>TGG</u>-3’, protospacer adjacent motif (PAM) is underlined) in the pKSE401 binary vector and carries a 1 bp insertion that causes a frameshift and a premature stop codon (Supplementary Fig. S1). The *GATA25*OX #3 and #13 lines were selected from independent transgenic lines following morphological assessment of 4-week-old plants and semi-quantitative reverse transcription PCR (RT-PCR) analysis of *GATA25* transcripts (Supplementary Fig. S2). The *gata25* mutant used in this study no longer carries the pKSE401-derived *zCas9* cassette, as confirmed by PCR with the pKSE401_F/R primers (Supplementary Fig. S3; Supplementary Table S1).

For soil cultivation, *Arabidopsis* seeds were cold-treated at 4 °C in darkness for 3 days and then sown in soil without surface sterilization. Plants were grown on soil in a growth chamber at 23 °C under LD (16 h light/8 h dark) or SD (8 h light/16 h dark) conditions with a light intensity of approximately 90–100 μmol m^-2^ s^-1^. For seedling analyses, seeds were surface-sterilized with 2% (w/v) sodium hypochlorite (NaClO) containing 0.05% (w/v) Triton X-100 for 5 min, followed by treatment with 70% ethanol for 10 sec, and then rinsed four times with sterile water on a clean bench. The sterilized seeds were sown on half-strength Murashige and Skoog (^1^/_2_ MS) medium supplemented with 1% (w/v) sucrose, 6 mM MES-KOH (pH 5.7), and 0.6% (w/v) phytoagar. After cold-treatment at 4 °C in darkness for 3 days, the seeds were grown in a growth chamber at 23 °C under LD or SD conditions with the same light intensity. *Nicotiana benthamiana* (*N. benthamiana*) plants were grown on soil at 23 °C under LD or SD conditions with the same light intensity and 5-week-old plants were used for agroinfiltration.

### Vector construction

To generate *GATA25*-overexpressing transgenic plants, the full-length coding sequence (CDS) of *GATA25* (AT4G24470) was amplified from leaf first-strand complementary DNA (cDNA) of *A. thaliana* by polymerase chain reaction (PCR) using gene-specific primers containing restriction enzyme sites (Supplementary Table S1). The amplified *GATA25* CDS was cloned into the laboratory-developed pTK-BMLC binary vector under control of the duplicated cauliflower mosaic virus (CaMV) 35S promoter (2×35S) (Kim et al., 2022). For dexamethasone (DEX)-inducible expression, the *GATA25* CDS was cloned into the laboratory-developed pTA7002-BMLC binary vector under control of the CaMV 2×35S promoter (Aoyama and Chua, 1997; Ko et al., 2009; Kim et al., 2022) (Fig. 2A). For epitope tagging, the *GATA25* CDS without the stop codon was amplified and fused in frame to the 5’ end of the monomeric red fluorescent protein (mRFP) sequence in the laboratory-developed pTK-mRFP-BMLC binary vector under control of the CaMV 2×35S promoter in this study (Supplementary Fig. S4A).

**Fig. 1.**
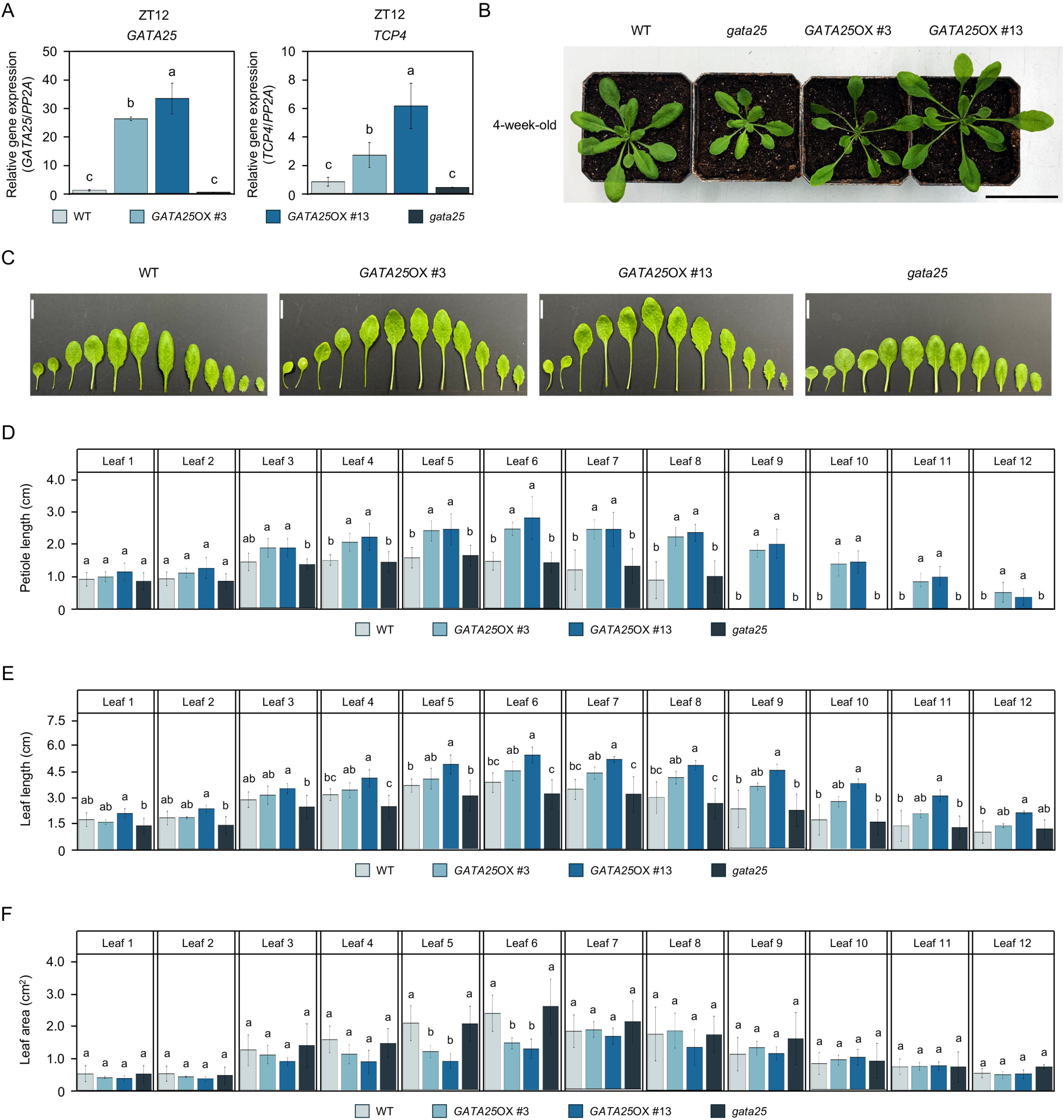
Functional characterization of GATA25 transcription factor in the regulation of *TCP4* expression and rosette leaf morphology from 4-week-old plants grown under LD conditions. (A) Transcript levels of *GATA25* and *TCP4* in leaves of WT, *GATA25*OX #3, *GATA25*OX #13, and *gata25* plants harvested at ZT12 under LD conditions. Transcript levels were normalized to *PP2A*. Data are presented as mean ± SD of three biological replicates. (B) Representative photographs of WT, *GATA25*OX #3, *GATA25*OX #13, and *gata25* plants. Scale bar, 7 cm. (C) Representative rosette leaves excised from WT, *GATA25*OX #3, *GATA25*OX #13, and *gata25* plants and arranged in order of emergence. Scale bar, 1 cm. (D-F) Petiole length (D), leaf length (E), and leaf blade area (F) of leaves 1 to 12 from WT, *GATA25*OX #3, *GATA25*OX #13, and *gata25* plants. The absence of bars in leaves 9 to 12 of WT and *gata25* indicates that no distinct petiole was formed. Data are presented as mean ± SD of six plants. Different letters indicate significant differences (one-way ANOVA with Tukey’s HSD test, *p* < 0.05).

**Fig. 2.**
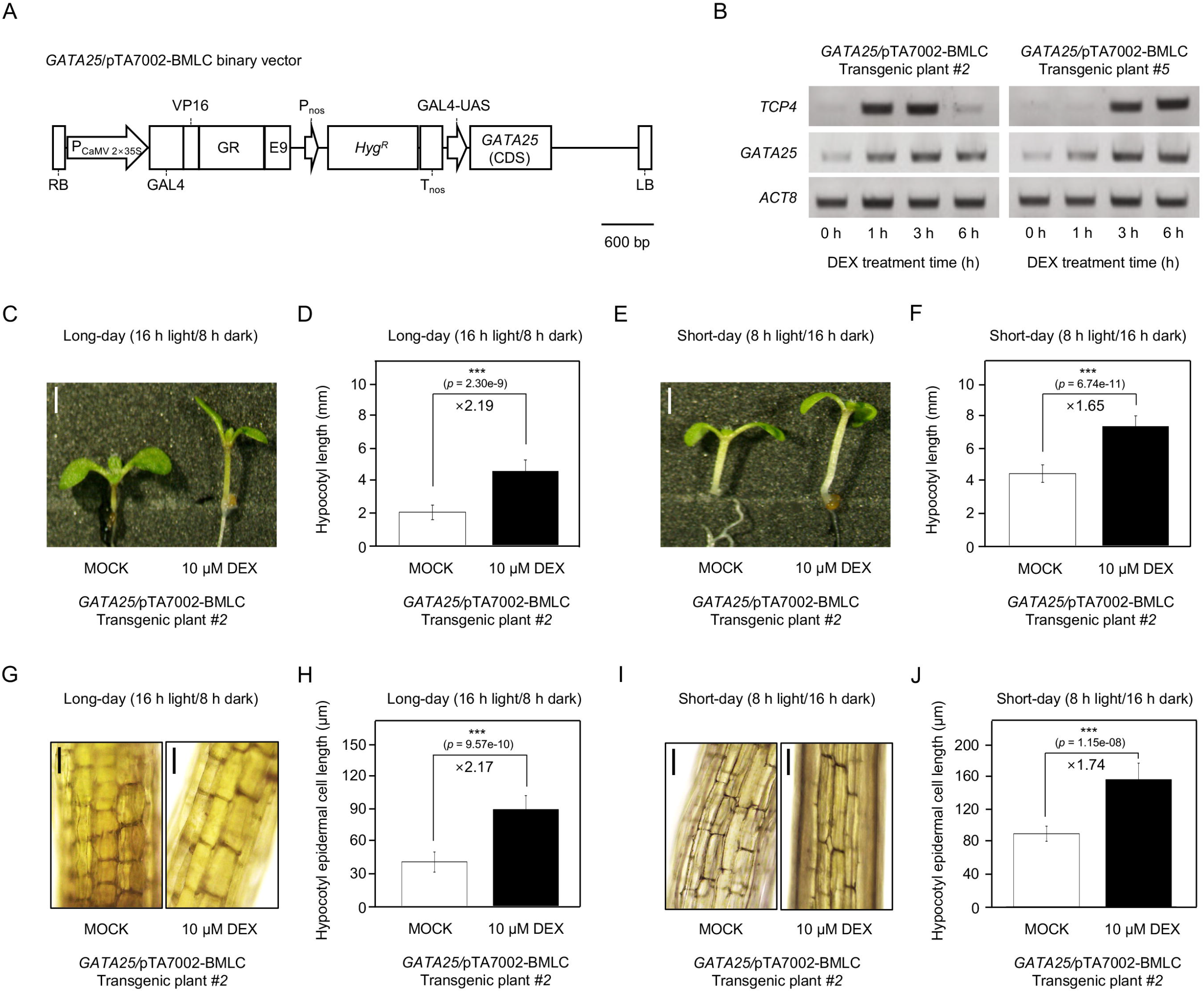
Inducible *GATA25* expression enhances *TCP4* expression and hypocotyl elongation in a photoperiod-dependent manner in 5-day-old seedlings. (A) Schematic diagram of the *GATA25*/pTA7002-BMLC binary vector for DEX-inducible expression. CDS, coding sequence; E9, pea rbsS-E9 transcriptional terminator; GAL4, yeast GAL4 DNA-binding domain; GAL4-UAS, GAL4 upstream activating sequence; GR, glucocorticoid receptor domain; *Hyg^R^*, hygromycin resistance gene; LB, left border; P_CaMV 2×35S_, cauliflower mosaic virus duplicated 35S promoter; P_nos_, nopaline synthase promoter; RB, right border; T_nos_, nopaline synthase terminator; VP16, herpes simplex virus VP16 transcriptional activation domain. Scale bar, 600 bp. (B) Semi-quantitative RT-PCR analysis of *GATA25* and *TCP4* transcripts in two independent inducible lines (#2 and #5) at 0, 1, 3, and 6 h after treatment with 10 μM DEX. *ACT8* was used as the internal control. (C, E) Representative photographs of seedlings of line #2 grown on ^1^/_2_ MS medium with or without 10 μM DEX under LD (C) or SD (E) conditions. Scale bar, 2 mm. (D, F) Hypocotyl length of seedlings under LD (D) and SD (F) conditions. Data are presented as mean ± SD of twelve seedlings. Numbers above the bars indicate the fold increase in DEX-treated seedlings relative to the corresponding mock-treated control. *p* values were determined by Welch’s *t*-test (\**p* < 0.05, \*\**p* < 0.005, and \*\*\**p* < 0.001). (G, I) Representative photographs of hypocotyl epidermal cells under LD (G) and SD (I) conditions. Scale bar, 50 μm. (H, J) Hypocotyl epidermal cell length under LD (H) and SD (J) conditions. Data are presented as mean ± SD of twelve seedlings, with twelve hypocotyl epidermal cells measured per seedling. Numbers above the bars indicate the fold increase in DEX-treated seedlings relative to the corresponding mock-treated control. *p* values were determined by Welch’s *t*-test (\**p* < 0.05, \*\**p* < 0.005, and \*\*\**p* < 0.001).

Plasmid DNA was extracted using an Exprep^TM^ Plasmid SV kit (GeneAll, Republic of Korea), and DNA fragments were purified using an Expin^TM^ Gel SV kit (GeneAll, Republic of Korea). Restriction enzymes were purchased from New England Biolabs (NEB, USA). *Escherichia coli* (*E. coli*) strain DH5α was used for cloning, and competent cells were prepared with 100 mM CaCl_2_ and 10% (v/v) glycerol and stored at -80 °C until use. All constructs were verified by Sanger sequencing (SolGent, Republic of Korea).

### Generation of transgenic plants

Each binary vector was introduced into *Agrobacterium tumefaciens* (*A. tumefaciens*) strain GV3101 by electroporation, and *A. thaliana* plants were transformed by the floral dip method (Clough and Bent, 1998; Zhang et al., 2006). Briefly, a single colony of *A. tumefaciens* carrying the construct was cultured in liquid Luria-Bertani (LB) medium supplemented with 10 μg mL^-1^ rifampicin, 30 μg mL^-1^ gentamicin, and 50 μg mL^-1^ kanamycin at 30 °C to a final optical density (OD) at 600 nm (OD_600nm_) of 0.8, and the cells were suspended in floral dip inoculation medium supplemented with 5% (w/v) sucrose and 0.05% (v/v) silwet L-77. Transformants carrying the pTK-BMLC or pTK-mRFP-BMLC construct were selected by spraying with Basta solution. Transformants carrying the pTA7002-BMLC construct were selected on ^1^/_2_ MS medium supplemented with 15 μg mL^-1^ hygromycin B and 0.6% (w/v) phytoagar as described previously (Harrison et al., 2006). Each selected transgenic line was advanced to the T_2_ generation, and the resulting plants were used for all analyses.

Transformants carrying the pKSE401 construct were selected on ^1^/_2_ MS medium supplemented with 50 μg mL^-1^ kanamycin and 0.6% (w/v) phytoagar as described previously (Harrison et al., 2006). The presence of the *zCas9* cassette of pKSE401 binary vector was confirmed by PCR using genomic DNA (gDNA) (Supplementary Fig. S3C). gDNA was extracted from young leaves by the modified Edwards method (Edwards et al., 1991; Kim et al., 2023a). Briefly, leaves were ground with a pestle, suspended in 300 μL of extraction solution [200 mM Tris-HCl (pH 9.0), 400 mM LiCl, 25 mM EDTA, and 1% (w/v) SDS], and vortexed. After centrifugation at 13,000 × g for 5 min, the supernatant was mixed with an equal volume of isopropanol and centrifuged at 13,000 × g for 10 min. The pellet was washed with 700 μL of 70% (v/v) ethanol, centrifuged at 13,000 × g for 5 min, and then dissolved in 20 μL of DNase-free distilled water. Each 20 μL PCR mixture contained 1 ng of gDNA, 0.2 mM of each primer, and EmeraldAmp^®^ GT PCR Master Mix (Takara Bio, Japan). The PCR conditions were as follows. Initial denaturation at 95 °C for 5 min, 34 cycles of denaturation at 95 °C for 30 sec, annealing at 55 °C for 30 sec, and extension at 72 °C for 1 min, followed by a final extension at 72 °C for 5 min. The products were separated on a 1% (w/v) agarose gel containing 1 μg mL^-1^ ethidium bromide (EtBr).

### DEX treatment of the inducible plants

Induction of *GATA25* in the DEX-inducible plants followed the GVG system protocol described previously (Aoyama and Chua, 1997; Ko et al., 2009; Kim et al., 2022). Two independent *GATA25*/pTA7002-BMLC lines (#2 and #5) were grown on soil for 2 weeks under LD conditions and sprayed with 10 μM DEX containing 0.02% (v/v) silwet L-77 until the rosettes were evenly wetted. Treated rosette leaves were harvested at 0, 1, 3, and 6 h after treatment, frozen immediately in liquid nitrogen, and stored at -80 °C until RNA extraction. Three plants were pooled per biological replicate, and three biological replicates were analyzed per time point.

For the hypocotyl length analysis, seeds of the *GATA25*/pTA7002-BMLC line #2 were surface-sterilized as described above and sown on ^1^/_2_ MS medium supplemented with 1% (w/v) sucrose, 6 mM MES-KOH (pH 5.7), 0.6% (w/v) phytoagar, and either 10 μM DEX or 0.01% (v/v) ethanol as a mock control. After cold-treatment at 4 °C in darkness for 3 days, the seeds were grown in a growth chamber at 23 °C under LD or SD conditions with the same light intensity described above.

### Phenotypic analysis

Rosette leaves were detached from 4-week-old plants grown under LD conditions and arranged sequentially according to their order of emergence on a white background. Petiole length, leaf length, and leaf blade area of leaves 1 to 12 were measured from digital images using ImageJ software version 1.54 (National Institutes of Health, USA). Six plants per line were analyzed for each of WT, *GATA25*OX #3, *GATA25*OX #13, and the *gata25* mutant, with each plant considered an independent biological replicate. Leaves were classified as lacking a distinct petiole when no clear constriction was observed between the leaf blade and the leaf base.

Hypocotyl length was measured in 5-day-old seedlings from images captured using a Leica EZ4E microscope (Leica, Germany) and analyzed with iSolution Lite software (IMT, Canada). Hypocotyl length was defined as the distance from the collet, where root hairs emerge, to the ‘v’ shaped junction between the cotyledons. Only seedlings that were not in contact with neighboring seedlings were included in the analysis, with 12 seedlings measured per treatment. Hypocotyl epidermal cell length was measured from images of the upper hypocotyl region captured using a Leica DM500 microscope (Leica, Germany). For each treatment, 12 epidermal cells were measured from each of 12 seedlings.

### RNA extraction, semi-quantitative RT-PCR, and qRT-PCR

Harvested tissues were ground to a fine powder with a pestle under liquid nitrogen. Total RNA was extracted using TRIzol reagent (Thermo Fisher Scientific, USA) according to the manufacturer’s instructions. Briefly, 1 mL of TRIzol reagent and 200 μL of chloroform/isoamyl alcohol (24:1, v/v) were added to the ground tissue, and the mixture was vortexed and centrifuged at 13,000 × g for 10 min at 4 °C. The supernatant (300 μL) was mixed with 300 μL of isopropanol, inverted, and centrifuged at 13,000 × g for 20 min at 4 °C. The pellet was washed with 700 μL of 70% (v/v) ethanol, centrifuged at 13,000 × g for 5 min at 4 °C, and then dissolved in 30 μL of RNase-free distilled water. Residual gDNA was removed by DNase I digestion using a RNeasy Plant Mini Kit (Qiagen, Germany). RNA concentration and purity were determined from the absorbance ratio at 260 and 280 nm using a NanoDrop ND-2000 spectrophotometer (Thermo Fisher Scientific, USA).

First-strand cDNA was synthesized from 2 μg of total RNA in a 20 μL reaction containing 1 μL of oligo (dT) primer (80 μM), 4 μL of 5× first-strand buffer, 1 μL of dNTP mixture (10 mM each), 2 μL of 0.1 M dithiothreitol (DTT), and 1 μL of SuperiorScript III reverse transcriptase (Enzynomics, Republic of Korea). Total RNA was denatured at 65 °C for 5 min before the reaction, reverse transcription was performed at 50 °C for 1 h in a T100^TM^ thermal cycler (Bio-Rad, USA), and the enzyme was inactivated at 70 °C for 15 min.

Semi-quantitative RT-PCR was performed on a T100^TM^ thermal cycler (Bio-Rad, USA) using EmeraldAmp^®^ GT PCR Master Mix (Takara Bio, Japan) in 20 μL reactions containing 2 μL of the synthesized first-strand cDNA and 0.2 mM of each primer. The primers used for RT-PCR are listed in Supplementary Table S2. The PCR conditions were as follows. Initial denaturation at 95 °C for 5 min, 30 cycles of denaturation at 95 °C for 30 sec, annealing at 55 °C for 30 sec, and extension at 72 °C for 30 sec, followed by a final extension at 72 °C for 5 min. *ACT8* (AT1G49240) was amplified as the internal control. The products were separated on a 1% (w/v) agarose gel containing 1 μg mL^-1^ ethidium bromide (EtBr). Band intensities were quantified using ImageJ software version 1.54 and normalized to those of *ACT8*.

Quantitative reverse transcription (qRT)-PCR was performed on a QuantStudio 3 Real-Time PCR System (Thermo Fisher Scientific, USA) using PowerUp^TM^ SYBR Green Master Mix (Thermo Fisher Scientific, USA) in 20 μL reactions containing 1 μL of the synthesized first-strand cDNA and 0.2 mM of each primer. The primers used for qRT-PCR are listed in Supplementary Table S2. Relative transcript levels were calculated by the 2^-ΔΔCt^ method (Livak and Schmittgen 2001) with *PP2A* (AT1G13320) as the reference gene. For the analysis of transcript levels during the dark period, leaves of 4-week-old plants grown under LD conditions were harvested at ZT12, ZT16, and ZT20. Each biological replicate consisted of leaves pooled from three plants, and three biological replicates were analyzed for each line and time point.

### Transcriptional activation analysis in *Arabidopsis* mesophyll protoplasts

The *TCP4* (AT3G15030) promoter region from -946 to +561 relative to the transcription start site (1,507 bp) was amplified from *A. thaliana* gDNA with primers containing *Pst*I and *Bam*HI restriction enzyme sites and cloned in front of the β*-glucuronidase* (*GUS*) gene of the pTrGUS vector, from which the CaMV 35S promoter had been removed, to generate the reporter construct (Fig. 3A). The *GATA25* CDS was cloned into the pTrTK-BMLC vector under the control of the CaMV 2×35S promoter (Kim et al., 2023b) to generate the effector (Fig. 3A). For the identification of direct targets, the *GATA25* CDS without the stop codon was fused in frame to the 5’ end of the glucocorticoid receptor domain (GR) sequence in the pTrTK-GR vector (Fig. 4A).

**Fig. 3.**
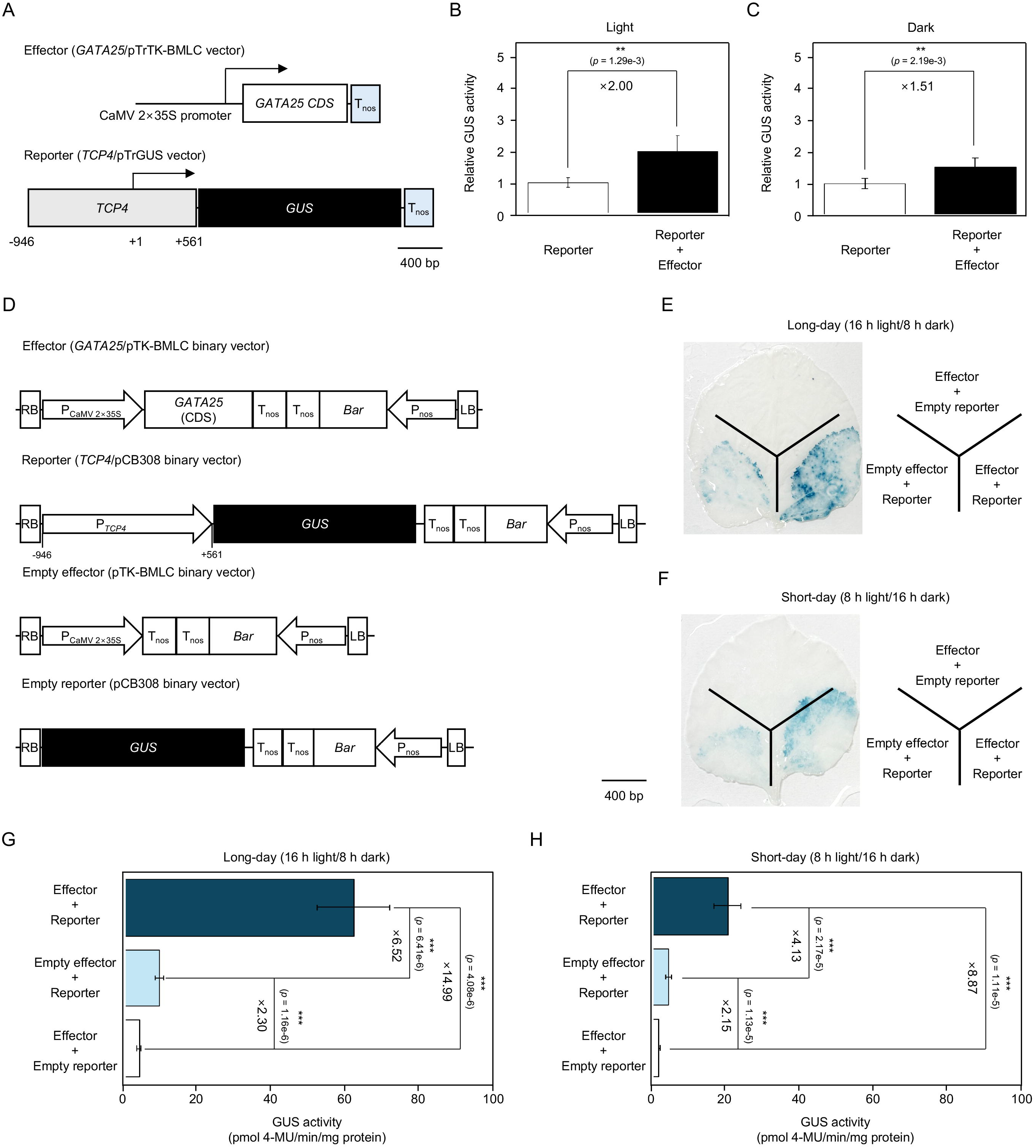
Transactivation of the *TCP4* promoter by GATA25 transcription factor under different light conditions. (A) Schematic diagrams of the *GATA25* effector and the *TCP4* promoter reporter vectors used in *Arabidopsis* mesophyll protoplasts. The effector vector contained the *GATA25* CDS driven by the CaMV 2×35S promoter. In the reporter vector, the *TCP4* promoter region from -946 to +561 relative to the transcription start site (+1) was fused to the *GUS* reporter gene. T_nos_, nopaline synthase terminator. Scale bar, 400 bp. (B, C) Relative GUS activity in *Arabidopsis* mesophyll protoplasts transfected with the *TCP4* promoter reporter alone or with both the *TCP4* promoter reporter and the *GATA25* effector. Protoplasts were subsequently incubated under continuous light (B) or continuous dark (C) conditions. (D) Schematic diagrams of effector, reporter, empty effector, and empty reporter binary vector used for agroinfiltration in *N. benthamiana* leaves. *Bar*, glufosinate/phosphinothricin resistance gene; LB, left border; P_CaMV 2×35S_, cauliflower mosaic virus duplicated 35S promoter; P_nos_, nopaline synthase promoter; RB, right border; T_nos_, nopaline synthase terminator. Scale bar, 400 bp. (E, F) Representative photographs of histochemical GUS staining of *N. benthamiana* leaves infiltrated with indicated combinations of vectors and subsequently grown under LD (E) and SD (F) conditions. (G, H) GUS activity in the infiltrated leaf region under LD (G) and SD (H) conditions. GUS activity was expressed in picomoles of 4-methylumbelliferone produced per minute per milligram of protein (pmol 4-MU/min/mg protein). Data are presented as mean ± SD of six biological replicates. *p* values were determined by Welch’s *t*-test (\**p* < 0.05, \*\**p* < 0.005, and \*\*\**p* < 0.001).

**Fig. 4.**
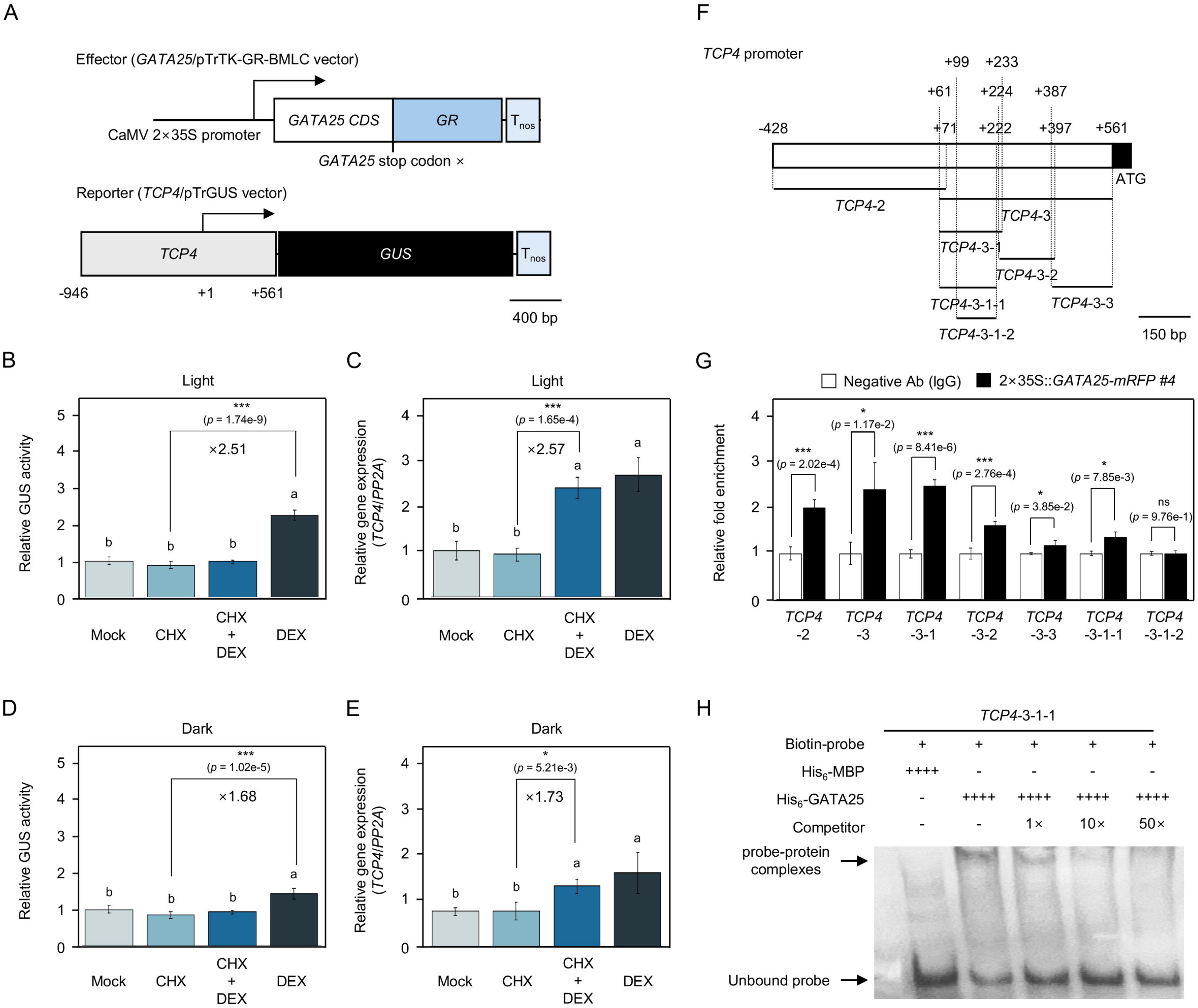
Direct binding of GATA25 transcription factor to the *TCP4* promoter. (A) Schematic diagrams of the *GATA25*-*GR* effector and *TCP4* promoter reporter vectors used in *Arabidopsis* mesophyll protoplasts. The effector vector contained the *GATA25* CDS without the stop codon, driven by the CaMV 2×35S promoter. In the reporter vector, the *TCP4* promoter region from -946 to +561 relative to the transcription start site (+1) was fused to the *GUS* reporter gene. GR, glucocorticoid receptor domain; T_nos_, nopaline synthase terminator. Scale bar, 400 bp. (B, D) Relative GUS activity in *Arabidopsis* mesophyll protoplasts co-transfected with the *GATA25*-*GR* effector and the *TCP4* promoter reporter and treated with Mock, 2 μM CHX, 10 μM DEX, or 2 μM CHX and 10 μM DEX under continuous light (B) or continuous dark (D) conditions. Data are presented as mean ± SD of three biological replicates. Different letters indicate significant differences (one-way ANOVA with Tukey’s HSD test, *p* < 0.05). *p* values were determined by Welch’s *t*-test (\**p* < 0.05, \*\**p* < 0.005, and \*\*\**p* < 0.001). (C, E) Endogenous *TCP4* transcript levels in the protoplasts transfected with the *GATA25-GR* effector alone and treated with Mock, 2 μM CHX, 10 μM DEX, or 2 μM CHX and 10 μM DEX under continuous light (C) or continuous dark (E) conditions. Transcript levels were normalized to *PP2A*. Data are presented as mean ± SD of three biological replicates. Different letters indicate significant differences (one-way ANOVA with Tukey’s HSD test, *p* < 0.05). *p* values were determined by Welch’s *t*-test (\**p* < 0.05, \*\**p* < 0.005, and \*\*\**p* < 0.001). (F) Schematic diagram of the *TCP4* promoter fragments used for ChIP-qPCR and EMSA. Numbers indicate positions relative to the transcription start site (+1). (G) ChIP-qPCR analysis of the *TCP4* promoter fragment in 2×35S::*GATA25*-*mRFP* #4 using anti-mRFP antibody. Enrichment is shown relative to the IgG control. Data are presented as mean ± SD of three biological replicates. *p* values were determined by Welch’s *t*-test (\**p* < 0.05, \*\**p* < 0.005, and \*\*\**p* < 0.001). (H) EMSA using the biotin-labeled *TCP4*-3-1-1 DNA probe, His_6_-GATA25 recombinant protein, and unlabeled *TCP4*-3-1-1 competitor at 1×, 10×, and 50× molar excess relative to the biotin-labeled *TCP4*-3-1-1 DNA probe. His_6_-MBP recombinant protein was used as the negative control. His_6_-MBP and His_6_-GATA25 recombinant protein were each used at concentration of 200 nM.

*Arabidopsis* mesophyll protoplasts were isolated from the leaves of 3-week-old *A. thaliana* and transfected by polyethylene glycol (PEG)-mediated transfection as described previously (Yoo et al., 2007). Briefly, 30 leaves were cut into 0.5 to 1 mm strips and transferred into 10 mL of enzyme solution [400 mM mannitol, 20 mM KCl, 20 mM MES-KOH (pH 5.7), 1.5% (w/v) cellulase, 0.1% (w/v) macerozyme, 10 mM CaCl_2_, and 0.1% (w/v) bovine serum albumin (BSA)], vacuum infiltrated for 30 min in a desiccator, and incubated for 2 h in darkness without shaking. The isolated protoplasts were diluted with 10 mL of W5 solution [5 mM KCl, 2 mM MES-KOH (pH 5.7), 125 mM CaCl_2_, and 154 mM NaCl], filtered through a nylon mesh, and allowed to settle on ice for 30 min. The protoplasts were resuspended in 2 mL of W5 solution, counted with a hemocytometer, and resuspended at 2 × 10^5^ mL-1 in MMG solution [400 mM mannitol, 15 mM MgCl_2_, and 4 mM MES-KOH (pH 5.7)]. For each transfection, 100 μL of protoplasts was mixed with 5 μg of the reporter, 5 μg of the effector or the empty vector, followed by the addition of 110 μL of PEG solution [30% (w/v) PEG 4000, 200 mM mannitol, and 100 mM CaCl_2_] for 15 min in darkness. The transfection was stopped with 440 μL of W5 solution, and the protoplasts were collected by centrifugation at 200 × g for 2 min, resuspended in 200 μL of WI solution [500 mM mannitol, 4 mM MES-KOH (pH 5.7), and 20 mM KCl], and incubated at room temperature for 12 h under continuous light (approximately 90–100 μmol m^-2^ s^-1^) or continuous darkness. Protoplasts transfected with the reporter alone were used as the control. For the DEX-inducible transcriptional activation analysis, 100 μL of protoplasts was mixed with 10 μg of the *GATA25*/pTrTK-GR effector, followed by the addition of 110 μL of PEG solution for 15 min in darkness. The transfection was stopped with 440 μL of W5 solution, and the protoplasts were collected by centrifugation at 200 × g for 2 min, resuspended in 200 μL of WI solution, and incubated for 2 h. Subsequently, the protoplasts were pretreated with 2 μM cycloheximide (CHX) for 30 min and subsequently treated with 10 μM DEX for 12 h. The protoplasts were lysed in 100 μL of lysis buffer [25 mM Tris-phosphate (pH 7.8), 1 mM DTT, 2 mM 1,2-diaminocyclohexane-N,N,N’,N’-tetraacetic acid, 10% (v/v) glycerol, and 1% (v/v) Triton X-100], incubated on ice for 5 min, and centrifuged at 13,000 × g for 2 min. For the GUS activity assay, 20 μL of the lysate was mixed with 100 μL of substrate solution [10 mM Tris-HCl (pH 8.0), 2 mM MgCl_2_, and 1 mM 4-methylumbelliferyl-β-D-glucuronide (MUG)] and incubated for 3 h at 37 °C, and the reaction was stopped with 880 μL of 200 mM Na_2_CO_3_. Fluorescence of the 4-methylumbelliferone (4-MU) product was measured on a SPECTRAmax GEMINI XS microplate spectrofluorometer (Molecular Devices, USA) at an excitation wavelength of 365 nm and an emission wavelength of 455 nm. Endogenous *TCP4* transcript levels in the DEX-treated protoplasts were analyzed by qRT-PCR as described above. Three biological replicates, each consisting of an independent protoplast preparation and transfection, were analyzed.

### Transactivation assay in *N. benthamiana*

The *TCP4* (AT3G15030) promoter region from -946 to +561 relative to the transcription start site (1,507 bp) was amplified from *A. thaliana* gDNA with primers containing *Xba*I and *Bam*HI restriction enzyme sites and cloned in front of the β*-glucuronidase* (*GUS*) gene of the pCB308 binary vector (Xiang et al., 1999) to generate the reporter (Fig. 3D). The *GATA25* CDS in the pTK-BMLC binary vector was used as the effector, and the empty pTK-BMLC and pCB308 vectors were used as the empty effector and reporter, respectively. Each binary vector was introduced into *A. tumefaciens* strain GV3101, and the cultures were resuspended in infiltration buffer [10 mM MES-KOH (pH 5.6), 10 mM MgCl_2_, and 150 μM acetosyringone] to a final OD_600nm_ of 0.8. Equal volumes of the effector and reporter cultures were mixed and infiltrated into different regions of the same leaf of *N. benthamiana* grown under LD conditions. The infiltrated plants were grown under LD or SD conditions for 3 days before the GUS activity analysis.

Histochemical GUS staining was performed as described previously (Jefferson et al., 1987; Kim et al., 2026). The infiltrated leaves were incubated overnight at 37 °C in GUS staining solution [50 mM NaPO_4_ (pH 7.0), 10 mM EDTA, 2 mM K_4_Fe(CN)_6_, 2 mM K_3_Fe(CN)_6_, 0.1% (v/v) Triton X-100, and 2.5 g L^-1^ X-gluc (5-bromo-4-chloro-3-indolyl-β-D-glucuronic acid)]. Chlorophyll and other pigments were removed by sequential incubation in increasing ethanol concentrations (25%, 50%, 75%, and 100%) at 37 °C, with each concentration applied overnight. The chlorophyll-free leaves were photographed using a Leica EZ4E microscope. For GUS activity assay, total protein was extracted from infiltrated leaves using extraction buffer [100 mM Tris-HCl (pH 8.5), 0.7 M sucrose, 0.1 M KCl, 50 mM EDTA, 1 mM phenylmethylsulfonyl fluoride (PMSF), 2% (v/v) β-mercaptoethanol, 10% (v/v) glycerol, 1% (v/v) Triton X-100, and protease inhibitors]. After centrifugation at 13,000 × g for 5 min at 4 °C, the supernatant was transferred to a new microcentrifuge tube, and the protein concentration was determined by the Bradford assay (Bradford 1976) using Bradford reagent (Bio-Rad, USA) with bovine serum albumin (BSA) as the standard (Bradford, 1976). An aliquot (20 μL) of the supernatant was mixed with 100 μL of substrate solution [10 mM Tris-HCl (pH 8.0), 2 mM MgCl_2_, and 1 mM 4-methylumbelliferyl-β-D-glucuronide (MUG)] and incubated for 3 h at 37 °C, and the reaction was stopped with 880 μL of 200 mM Na_2_CO_3_. GUS activity was determined by measuring 4-MU production from MUG on a SPECTRAmax GEMINI XS microplate spectrofluorometer as described above. GUS activity was expressed as pmol of 4-MU per min per milligram of protein (pmol 4-MU/min/mg protein).

### Chromatin immunoprecipitation (ChIP)-qPCR

The 4-week-old 2×35S::*GATA25*-*mRFP* #4 plants grown under LD conditions were harvested at ZT12 and vacuum-infiltrated for 30 min in 100 mL of phosphate-buffered saline (PBS) [3 mM Na_2_HPO_4_, 3 mM NaH_2_PO_4_ (pH 7.0), and 130 mM NaCl] containing 2 mM disuccinimidyl glutarate. After a wash with PBS, the tissue was vacuum-infiltrated for 20 min in 100 mL of PBS containing 1% (v/v) formaldehyde, and cross-linking was quenched with 125 mM glycine under vacuum for 5 min. The tissue was rinsed twice with deionized water, frozen in liquid nitrogen, and stored at -80 °C until use. Nuclei were extracted with three extraction buffers, each containing 10 mM Tris-HCl (pH 8.0), 5 mM β-mercaptoethanol, 1× protease inhibitor cocktail, and 1 mM PMSF, together with 0.4 M sucrose and 10 mM MgCl_2_ (extraction buffer 1), 0.25 M sucrose, 10 mM MgCl_2_, and 1% (v/v) Triton X-100 (extraction buffer 2), or 1.7 M sucrose, 2 mM MgCl_2_, and 0.15% (v/v) Triton X-100 (extraction buffer 3). The tissue was ground to a fine powder in liquid nitrogen, suspended in 10 mL of extraction buffer 1, filtered through Miracloth (Merck Millipore, Germany), and centrifuged at 5,000 × g for 10 min at 4 °C. The pellet was resuspended in 1 mL of extraction buffer 2 and centrifuged under the same conditions. The resulting pellet was resuspended in 300 μL of extraction buffer 3, layered on top of 600 μL of extraction buffer 3, and centrifuged at 15,000 × g for 60 min at 4 °C. The nuclear pellet was lysed in 300 μL of nuclei lysis buffer [50 mM Tris-HCl (pH 8.0), 10 mM EDTA, 1% (w/v) SDS, 1× protease inhibitor cocktail, and 1 mM PMSF], and the chromatin was sheared with a Bioruptor (Diagenode, Belgium) at 4 °C to an average fragment size of approximately 500 bp. For each preparation, 150 μL of the sheared chromatin was used, and 15 μL was stored at -20 °C as the input. The remaining chromatin was diluted tenfold with ChIP dilution buffer [16.7 mM Tris-HCl (pH 8.0), 167 mM NaCl, 1.2 mM EDTA, 1.1% (v/v) Triton X-100, and 1 mM PMSF] and divided into two portions. One portion was incubated with anti-mRFP monoclonal antibody-conjugated magnetic beads (MBL, Japan), and the other was incubated with normal mouse IgG and Dynabeads Protein G (Invitrogen, USA) as the negative control, overnight at 4 °C. The beads were washed at 4 °C for 5 min per wash in 1 mL of low-salt wash buffer [20 mM Tris-HCl (pH 8.0), 150 mM NaCl, 2 mM EDTA, 0.1% (w/v) SDS, and 1% (v/v) Triton X-100] twice, high-salt wash buffer (low-salt wash buffer containing 500 mM NaCl) three times, LiCl wash buffer [10 mM Tris-HCl (pH 8.0), 0.25 mM LiCl, 1 mM EDTA, 1% (v/v) NP-40, and 1% (w/v) sodium deoxycholate] once, and TE buffer [10 mM Tris-HCl (pH 8.0) and 1 mM EDTA] twice. DNA was recovered from the beads and the input samples with 200 μL of 10% (w/v) Chelex^®^ 100 Resin (Bio-Rad, USA) by heating at 98 °C for 10 min, digestion with 2 μL of proteinase K (10 mg mL^-1^) at 37 °C for 20 min, and heating again at 98 °C for 10 min, and the supernatant was collected after centrifugation at 15,000 × g for 5 min at 4 °C.

qPCR was performed as described above with the ChIP DNA and tenfold-diluted input DNA as templates using the primers listed in Supplementary Table S2. The relative fold enrichment was calculated by the comparative Ct method. For each fragment, ΔCt was obtained by subtracting Ct of the IgG control samples from the Ct of each sample, and the 2^-ΔCt^ value of each sample was divided by the 2^-ΔCt^ value of the IgG control samples, so that the IgG control was set to 1. Three biological replicates, each from an independent chromatin preparation, were analyzed.

### Recombinant protein purification and electrophoretic mobility shift assay (EMSA)

The *GATA25* CDS was cloned into pET28a (+) vector (Novagen, USA), and the *maltose binding protein (MBP)* sequence was cloned into pET21a (+) (Novagen, USA) as the control (Supplementary Table S1). The pET28a-*GATA25* and pET21a-*MBP* constructs were separately transformed into *E. coli* strain BL21 (DE3). The transformed cells were cultured in LB medium supplemented with 50 μg mL^-1^ kanamycin for pET28a-*GATA25* or 100 μg mL^-1^ ampicillin for pET21a-*MBP* at 37 °C until the OD_600nm_ reached 0.5. Protein expression was induced with 0.5 mM isopropyl β-D-thiogalactopyranoside (IPTG) at 18 °C overnight. Cells were harvested, resuspended in binding buffer [500 mM NaCl, 20 mM Tris-HCl (pH 7.9), and 5 mM imidazole], and lysed by sonication. Cell debris was removed by centrifugation at 15,000 × g for 20 min at 4 °C, and the His_6_-tagged recombinant proteins were purified from the cleared lysate by Ni^2+^-NTA affinity chromatography (Qiagen, USA) according to the instructions of the manufacturer and dialyzed against binding buffer overnight at 4 °C (Kim et al., 2024). Protein purity was confirmed by 12% SDS-polyacrylamide gel electrophoresis (PAGE) (Supplementary Fig. S5A), and protein concentration was determined by the Bradford assay.

For EMSA using unlabeled DNA, three *TCP4* promoter fragments (*TCP4*-1, *TCP4*-2, and *TCP4*-3) were amplified from *A. thaliana* gDNA using the primers listed in Supplementary Table S3. Each fragment was incubated with increasing concentrations of His_6_-GATA25 recombinant protein (0, 50, 75, 100, or 200 nM) or 200 nM His_6_-MBP recombinant protein as a negative control for 30 min at room temperature. The DNA-protein complexes were then separated on a 0.6% (w/v) agarose gel. For EMSA using biotin-labeled DNA probes, the *TCP4*-3-1-1 and *TCP4*-3-1-2 fragments were labeled at their 3’ ends with biotin-11-UTP using terminal deoxynucleotidyl transferase and a Biotin 3’ End DNA Labeling Kit (Thermo Fisher Scientific, USA). The labeled probes were subsequently purified by chloroform/isoamyl alcohol extraction. Each binding reaction (20 μL) contained 20 fmol of biotin-labeled DNA probe, His_6_-GATA25 recombinant protein (0, 75, 100, or 200 nM) or His_6_-MBP recombinant protein (200 nM) as a negative control, and binding buffer [10 mM Tris-HCl (pH 7.5), 50 mM KCl, 1 mM Dithiothreitol (DTT), 2.5% (v/v) glycerol, 5 mM MgCl_2_, 0.05% (v/v) NP-40, and 50 ng μL^-1^ poly(dI·dC)]. The reaction mixtures were incubated for 30 min at room temperature. For competition assays, unlabeled *TCP4*-3-1-1 fragments were added at 1-, 10-, or 50-fold molar excess relative to the labeled probe. The probe-protein complexes were separated by electrophoresis on a 6% native polyacrylamide gel, transferred to a nylon membrane, and detected by chemiluminescence using X-ray films.

### Yeast two-hybrid (Y2H) assay

The CDSs of *GATA25* and *COP1* (AT2G32950) were cloned into pGADT7 and pGBKT7 vectors (Clontech, USA) in reciprocal combinations (Supplementary Table S1). The resulting prey and bait constructs were co-transformed into *Saccharomyces cerevisiae* strain AH109 by the lithium acetate (LiAc) method (Gietz and Schiestl, 2007; Mehla et al., 2015). Transformants were selected on synthetic dropout (SD) medium lacking leucine and tryptophan (SD/-Leu/-Trp). To assess protein-protein interactions, yeast cells were harvested from 10 mL of culture at an OD_600nm_ of 0.8 and subjected to 10-fold serial dilutions (10^0^–10^-5^) in Phosphate-buffered saline (PBS). Aliquots of each dilution were spotted onto YEPAD medium as a growth control, SD/-Leu/-Trp medium, SD medium lacking leucine, tryptophan, histidine, and adenine (SD/-Leu/-Trp/-His/-Ade), and SD/-Leu/-Trp/-His/-Ade medium supplemented with 40 μg mL^-1^ X-α-Gal. Yeast growth and reporter activity were evaluated after incubation at 30 °C for 3 days. Co-transformation of each construct with the corresponding empty vector served as a negative control.

### Bimolecular fluorescence complementation (BiFC) assay

The CDSs of *GATA25* and *COP1* without stop codons were cloned into pVenus^N173^ and pVenus^C155^ vectors in reciprocal combinations (Supplementary Table S1). Each construct was transformed into *A. tumefaciens* strain GV3101 and equal volumes of the cultures carrying the respective constructs were mixed and co-infiltrated into leaves of *N. benthamiana* using infiltration buffer containing 50 μM MG132 to inhibit proteasomal degradation. Full-length Venus was used as a positive control, whereas the combination of empty pVenus^N173^ and pVenus^C155^ vectors served as a negative control. The infiltrated plants were incubated in darkness for 3 days, and Venus fluorescence in the epidermal cells of the infiltrated regions was observed with a BX53 fluorescence microscope (Olympus, Japan) equipped with an EYFP/Venus/Citrine fluorescence filter set (Chroma Technology, USA) with excitation and emission wavelength ranges of 485–505 nm and 525–555 nm, respectively.

### Statistical analysis

Data are presented as the mean ± standard deviation (SD). Comparisons among three or more groups were performed using one-way analysis of variance (ANOVA) followed by Tukey’s honestly significant difference (HSD) test, with different letters indicating significant differences at *p* < 0.05. Comparisons between two groups were performed using a two-tailed unpaired Welch’s *t*-test (Zimmerman, 2004; Ruxton, 2006; Delacre et al., 2017). Statistical significance was indicated by \**p* < 0.05, \*\**p* < 0.005, and \*\*\**p* < 0.001. All statistical analyses were performed using SAS software version 9.4 (SAS Institute, USA). iological replicates consisted of independent plants, independent protoplast preparations, or independent infiltrated leaf regions, as specified in the corresponding experimental sections. The number of biological replicates is indicated in each figure legend.

## Results

### GATA25 transcription factor promotes petiole elongation and *TCP4* expression in rosette leaves

To investigate whether the GATA25 transcription factor affects leaf development, we examined the rosette leaves of two independent *GATA25*-overexpressing lines (*GATA25*OX #3 and #13) and the *gata25* mutant grown under LD conditions for 4 weeks (Fig. 1; Supplementary Fig. S1-S3). At ZT12, *GATA25* transcript levels were higher in *GATA25*OX #3 and #13 than in WT, with the highest level in *GATA25*OX #13. Similarly, *TCP4* transcript levels were higher in both *GATA25*OX lines than in WT, with the highest level in *GATA25*OX #13, whereas no significant difference was observed between the *gata25* mutant and WT (Fig. 1A). Both *GATA25*OX lines showed longer petioles than WT and the *gata25* mutant from leaf 4 to leaf 8, whereas no significant difference in petiole length was detected between WT and the *gata2*5 mutant in any leaf (Fig. 1B-D). Notably, distinct petioles were observed in leaves 9 to 12 of both *GATA25*OX lines but not in those of WT or the *gata25* mutant (Fig. 1C, D). Leaf length was greater in *GATA25*OX #13 than in WT from leaf 4 to leaf 12, while *GATA25*OX #3 showed intermediate values between WT and *GATA25*OX #13 (Fig. 1E). These differences between two independent *GATA25*OX lines were consistent with their relative *GATA25* and *TCP4* transcript levels (Fig. 1A). Interestingly, leaf 5 and leaf 6 of both *GATA25*OX lines had significantly smaller areas than those of WT and the *gata25* mutant, whereas no significant difference in leaf area was observed in the other leaves (Fig. 1F). This reduction in leaf area is consistent with previous studies showing that increased TCP4 activity restricts leaf growth by limiting cell proliferation (Sarvepalli and Nath, 2011; Challa et al., 2019). These results suggest that elevated *TCP4* expression may contribute to the petiole elongation and reduced leaf area associated with *GATA25* overexpression.

### Inducible *GATA25* expression increases *TCP4* expression and hypocotyl elongation under different photoperiod conditions

Given the potential involvement of *TCP4* in *GATA25*-mediated leaf development, we further investigated whether *GATA25* regulates hypocotyl elongation through *TCP4*, based on a previous study demonstrating that *TCP4* promotes hypocotyl elongation in a light-dependent manner, as *TCP4* induction failed to enhance hypocotyl elongation in dark-grown seedlings (Challa et al., 2016). We hypothesized that *GATA25* may promote hypocotyl elongation in association with *TCP4* expression and that this effect may be greater under light than dark conditions. To test this hypothesis, we first examined the temporal relationship between *GATA25* induction and *TCP4* expression using two independent DEX-inducible *GATA25*/pTA7002-BMLC transgenic lines (#2 and #5) (Fig. 2A). In DEX-inducible transgenic line #2, *GATA25* transcript levels increased within 1 h of treatment with 10 μM DEX and remained elevated for up to 6 h, and *TCP4* transcript levels increased in response to *GATA25* induction at 1 h and 3 h before declining at 6 h (Fig. 2B). In contrast, in DEX-inducible transgenic line #5, *GATA25* transcript levels increased at 1 h, whereas *TCP4* transcript levels did not increase in response to *GATA25* induction until 3 h (Fig. 2B). These results indicate that *GATA25* induction and the subsequent transcriptional activation of *TCP4* occurred earlier in DEX-inducible transgenic line #2 than in line #5. Therefore, DEX-inducible transgenic line #2 was selected for subsequent experiments to investigate *GATA25*-mediated *TCP4* transcriptional regulation and hypocotyl elongation following DEX treatment.

In DEX-inducible transgenic line #2, treatment with 10 μM DEX increased hypocotyl length by 2.19-fold relative to the mock-treated control under LD conditions and by 1.65-fold under SD conditions (Fig. 2C-F). Similarly, hypocotyl epidermal cell length increased by 2.17-fold under LD conditions and by 1.74-fold under SD conditions (Fig. 2G-J). These fold increases in cell length were comparable to those in hypocotyl length, indicating that the hypocotyl elongation induced by *GATA25* resulted mainly from cell elongation. Interestingly, both mock-treated and DEX-treated seedlings had longer hypocotyls under SD conditions than under LD conditions, whereas the relative increase induced by *GATA25* was larger under LD conditions (Fig. 2D, F, H, J). These results suggest that the GATA25 transcription factor promotes hypocotyl cell elongation together with *TCP4* expression and that its relative contribution is greater under LD conditions.

### GATA25 transcription factor transactivates the *TCP4* promoter, with higher activity under light and LD conditions

To test whether the GATA25 transcription factor activates the *TCP4* promoter, we co-transfected *Arabidopsis* mesophyll protoplasts with the *TCP4* promoter (-946 to +561)-driven GUS reporter and the CaMV 2×35S promoter-driven *GATA25* effector (Fig. 3A). The *GATA25* effector increased GUS activity by 2.00-fold relative to the reporter alone under continuous light and by 1.51-fold under continuous dark conditions (Fig. 3B, C). In *N. benthamiana* leaves grown under LD or SD conditions for 3 days after agroinfiltration, GUS staining was strongest in the region co-infiltrated with the effector and reporter, weaker in the region with the empty effector and reporter, and barely detected in the region with the effector and empty reporter (Fig. 3D-F). Under LD conditions, GUS activity with the effector and reporter was 6.52-fold higher than that with the empty effector and reporter and 14.99-fold higher than that with the effector and empty reporter (Fig. 3G). Under SD conditions, the corresponding increases were 4.13-fold and 8.87-fold, respectively (Fig. 3H). The empty effector and reporter combination also showed 2.30-fold and 2.15-fold higher GUS activity than the effector and empty reporter combination under LD and SD conditions, respectively, indicating basal *TCP4* promoter activity in *N. benthamiana* leaves (Fig. 3G, H). Notably, GUS activity in the region co-infiltrated with the effector and reporter was 2.98-fold higher under LD conditions than under SD conditions (Supplementary Fig. S6), and the fold activation by GATA25 transcription factor was also larger under LD conditions (Fig. 3G, H). These results suggest that the GATA25 transcription factor activates the *TCP4* promoter and that this activation is enhanced under light and LD conditions.

### GATA25 transcription factor directly binds to the *TCP4* promoter

To determine whether *TCP4* is a direct target of the GATA25 transcription factor, we used in the GATA25-GR system in *Arabidopsis* mesophyll protoplasts (Fig. 4A). The GR-based inducible system enables the DEX-dependent activation of GR-fused transcription factors through their nuclear translocation without requiring *de novo* protein synthesis, allowing the use of CHX, a protein synthesis inhibitor, to distinguish primary transcriptional responses from those requiring *de novo* synthesized proteins (Yamaguchi et al., 2015; Shimada et al., 2022). Under continuous light conditions, treatment with DEX increased GUS activity by 2.51-fold relative to CHX treatment alone, whereas GUS activity with CHX and DEX remained at the mock level (Fig. 4B). Similarly, under continuous dark conditions, treatment with DEX increased GUS activity by 1.68-fold relative to CHX treatment alone, whereas GUS activity with CHX and DEX remained at the mock level (Fig. 4D). The suppression of DEX-induced GUS activity by CHX is consistent with the inhibition of *de novo* GUS protein synthesis, despite the expected DEX-dependent activation of GATA25-GR. Notably, GUS activity following DEX treatment was 1.55-fold higher under continuous light than under continuous dark conditions (Supplementary Fig. S7). Together, these results indicate that the GATA25-GR system exhibited the expected DEX-responsive reporter activity under both light and dark conditions, with a greater relative increase observed under light conditions, supporting its applicability for investigating *GATA25*-mediated transcriptional regulation. To determine whether GATA25 transcription factor can activate *TCP4* transcription independently of *de novo* protein synthesis, we examined endogenous *TCP4* transcript levels under the same treatment conditions. Under continuous light conditions, co-treatment with CHX and DEX increased endogenous *TCP4* transcript levels by 2.57-fold relative to CHX treatment alone, reaching a level comparable to that observed with DEX treatment alone (Fig. 4C). Under continuous dark conditions, co-treatment with CHX and DEX increased endogenous *TCP4* transcript levels by 1.73-fold relative to CHX treatment alone (Fig. 4E). The accumulation of endogenous *TCP4* transcripts following DEX treatment in the presence of CHX suggests that GATA25 transcription factor can activate *TCP4* transcription independently of *de novo* protein synthesis. Notably, the greater fold increase in *TCP4* transcript levels under continuous light than under continuous dark conditions suggests that *GATA25*-mediated transcriptional activation of *TCP4* may be enhanced under light conditions.

To further investigate whether GATA25 transcription factor directly regulates *TCP4* transcription, we performed ChIP-qPCR using 2×35S::*GATA25*-*mRFP* #4 transgenic lines, which exhibited higher *GATA25* transcript levels and longer hypocotyls than WT plants (Fig. S4). Compared with the IgG control, significant enrichment was detected in the *TCP4*-2 (-428 to +71), *TCP4*-3 (+61 to +561), *TCP4*-3-1 (+61 to +233), *TCP4*-3-2 (+224 to +397), *TCP4*-3-3 (+387 to +561), and *TCP4*-3-1-1 (+61 to +222) regions. In contrast, no significant enrichment was detected in the *TCP4*-3-1-2 region (+99 to +222) (Fig. 4F, G).

To determine whether GATA25 transcription factor directly binds to these regions, we initially screened three *TCP4* promoter DNA fragments (*TCP4*-1, *TCP4*-2, and *TCP4*-3) using EMSA. His_6_-GATA25 recombinant protein formed DNA-protein complexes with the *TCP4*-1 (-946 to -415), *TCP4*-2 (-428 to +71), and *TCP4*-3 (+61 to +561) fragments at concentrations of 100 and 200 nM, whereas His_6_-MBP recombinant protein did not (Supplementary Fig. S5A-C). Based on the ChIP-qPCR results showing significant enrichment in the *TCP4*-3-1-1 region but not in the *TCP4*-3-1-2 region (Fig. 4G), we further examined these two overlapping fragments to narrow down the GATA25-binding region. His_6_-GATA25 recombinant protein bound to the biotin-labeled *TCP4*-3-1-1 probe at a concentration of 100 nM or higher, accompanied by a decrease in the unbound probe, whereas no binding was observed with the *TCP4*-3-1-2 probe (Supplementary Fig. S5D-F). Furthermore, the His_6_-GATA25 recombinant protein-*TCP4*-3-1-1 complex decreased in the presence of a 1× unlabeled competitor and was barely detectable with 10× and 50× unlabeled competitors, accompanied by a corresponding increase in the unbound probe (Fig. 4H). These results suggest that the +61 to +98 region of *TCP4* promoter, which is present in *TCP4*-3-1-1 but absent from *TCP4*-3-1-2, contains a specific DNA sequence recognized by GATA25 transcription factor. This region lacks the canonical GATA transcription factor-binding motif (Omichinski et al., 1993; Ravagnani et al., 1997; Starich et al., 1998; Maxon and Herskowitz, 2001; Froehlich et al., 2002; Teakle et al., 2002; Reyes et al., 2004; Behringer and Schwechheimer, 2015; Xu et al., 2017), but contains two 5’-NATC-3’ motifs and one 5’-ATCA(A/G)-3’ motif, corresponding to 5’- GATN-3’ and 5’-(C/T)TGAT-3’ motifs on the antisense strand, respectively (Fig. S8). Taken together, these results suggest that GATA25 transcription factor directly regulates *TCP4* transcription by binding to a region containing putative GATA-binding motifs within the *TCP4* promoter.

### GATA25 transcription factor interacts with COP1, suggesting a potential role in the dark-dependent regulation of *TCP4*

To investigate whether GATA25 activity is reduced during the dark period, we focused on the E3 ubiquitin ligase COP1. COP1 is known to negatively regulate light signaling by promoting the ubiquitination and subsequent proteasomal degradation of various transcription factors, including HY5, particularly in dark conditions (Osterlund et al., 2000; Holm et al., 2002; Hoecker, 2017). In addition, COP1 has been shown to interact with CO, a transcriptional regulator containing a CCT domain, and promote its degradation in dark conditions (Jang et al., 2008; Liu et al., 2008). Given the role of COP1 in the post-translational regulation of transcription factors, we investigated whether GATA25 physically interacts with COP1.

To determine whether the GATA25 transcription factor interacts with the COP1, we performed Y2H assays (Fig. 5A). Yeast cells co-expressing GATA25 fused to the GAL4 activation domain (pGADT7-GATA25) and COP1 fused to the GAL4 DNA-binding domain (pGBKT7-COP1) grew on SD/-Leu/-Trp/-His/-Ade medium at dilutions up to 10^-5^ and developed a blue color in the presence of X-α-Gal. Similar results were observed with the GATA25 fused to the GAL4 DNA-binding domain (pGBKT7-GATA25) and COP1 fused to the GAL4 activation domain (pGADT7-COP1) combination, whereas all negative control combinations grew only on SD/- Leu/-Trp medium. Consistent with these findings, BiFC assays in *N. benthamiana* leaves incubated in dark conditions in the presence of 50 μM MG132 revealed Venus fluorescence in both GATA25-Venus^N173^/COP1-Venus^C155^ and COP1-Venus^N173^/GATA25-Venus^C155^ combinations in the nucleus (Fig. 5B). In contrast, the Venus positive control exhibited fluorescence throughout the cell, including the nucleus, whereas no fluorescence was detected in the Venus^N173^/Venus^C155^ combination used as the negative control.

**Fig. 5.**
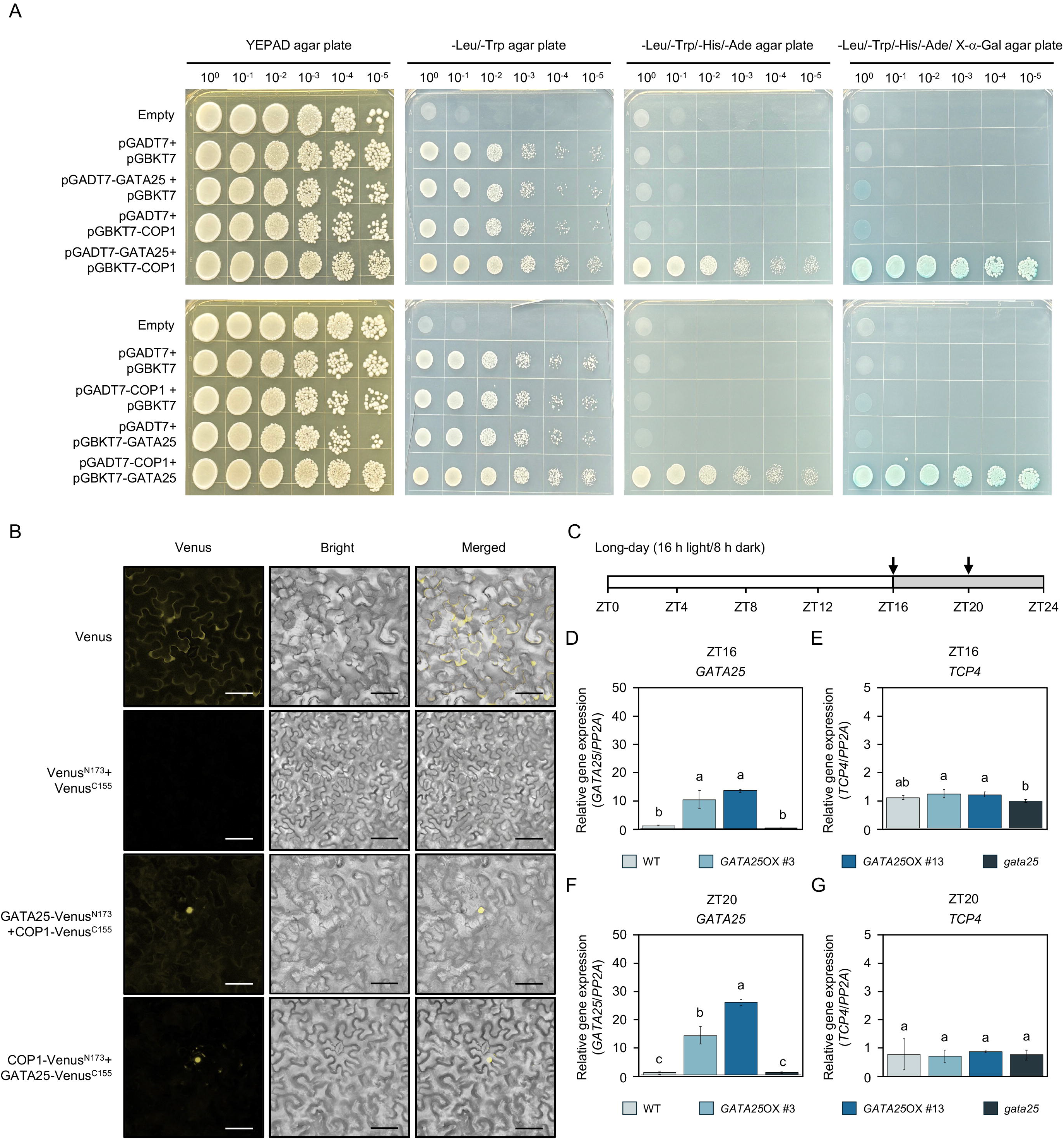
Molecular mechanisms underlying the physical interaction between GATA25 and COP1 in regulation of *TCP4*. (A) Y2H assay showing interaction between GATA25 and COP1. AH109 cells co-transformed with the indicated combination of pGADT7 and pGBKT7 vectors were subjected to 10-fold serial dilutions (10^0^ to 10^-5^) and spotted onto YEPAD, SD/-Leu/-Trp, SD/-Leu/-Trp/-His/- Ade, and SD/-Leu/-Trp/-His/-Ade agar plates supplemented with 40 μg mL^-1^ X-α-Gal. Co-transformation with the corresponding empty vectors was used as the negative control. (B) BiFC assay analysis detecting GATA25-COP1 interaction in *N. benthamiana* leaves. Full-length Venus and the combination of Venus^N173^ and Venus^C155^ were used as the positive and negative controls, respectively. Scale bars, 50 μm. (C) Schematic diagram of the LD photoperiod and sampling times. Black arrows indicate the time points. White box indicates the light period, and gray box indicates the dark period. (D-G) Transcript levels of *GATA25* (D, F) and *TCP4* (E, G) in leaves of 4-week-old WT, *GATA25*OX #3, *GATA25*OX #13, and *gata25* plants harvested at ZT16 (D, E) and ZT20 (F, G). Transcript levels were normalized to *PP2A*. Data are presented as mean ± SD of three biological replicates. Different letters indicate significant differences (one-way ANOVA with Tukey’s HSD test, *p* < 0.05).

Furthermore, we examined whether *GATA25* overexpression affects *TCP4* expression during the dark period. Transcript levels were analyzed in leaves of 4-week-old plants at ZT16 and ZT20, both of which fall within the dark period under LD conditions (Fig. 5C). *GATA25* transcript levels remained significantly higher in both *GATA25*OX lines than in WT plants at both time points (Fig. 5D, F). However, despite the elevated *GATA25* transcript levels, *TCP4* transcript levels in the *GATA25*OX lines were not significantly different from those in WT plants at either ZT16 or ZT20 (Fig. 5E, G). This contrasts with the increased *TCP4* transcript levels observed in the *GATA25*OX lines at ZT12 (Fig. 1A). At ZT16, *TCP4* transcript levels in the *gata25* mutant were significantly lower than those in the *GATA25*OX lines but were not significantly different from those in WT plants. At ZT20, no significant differences in *TCP4* transcript levels were detected among WT plants, the *GATA25*OX lines, and the *gata25* mutant (Fig. 5E, G). Together, these results indicate that *GATA25* overexpression does not enhance *TCP4* expression during the dark period, despite the sustained accumulation of *GATA25* transcripts. This suggests that the ability of GATA25 transcription factor to promote *TCP4* expression may be attenuated in dark conditions, potentially through its interaction with COP1.

## Discussion

This study elucidates the role of the GATA25 transcription factor in regulating light-dependent cell elongation through the direct transcriptional activation of *TCP4*. Beyond identifying GATA25 as a positive regulator of petiole and hypocotyl elongation, our findings reveal that the *GATA25*-mediated activation of *TCP4* is enhanced under light and LD conditions and that GATA25 physically interacts with the E3 ubiquitin ligase COP1. Together, these findings suggest a potential regulatory link between light signaling, transcriptional regulation, and cell elongation, providing new insights into how plants coordinate growth with daily light/dark cycles.

Our findings identify GATA25 as a positive regulator of petiole and hypocotyl elongation, potentially acting through the transcriptional activation of *TCP4*. Both independent *GATA25*OX lines exhibited longer petioles and higher *TCP4* transcript levels than WT plants, with *GATA25*OX #13 showing the highest *GATA25* and *TCP4* transcript levels and the greatest leaf length (Fig. 1). This positive association between *GATA25* expression, *TCP4* transcript abundance, and elongation phenotypes suggests that the transcriptional regulation of *TCP4* contributes to *GATA25*-mediated growth. Consistent with this interpretation, inducible expression of *GATA25* rapidly increased *TCP4* transcript levels and promoted hypocotyl elongation primarily through increased cell elongation (Fig.c2). Previous studies have demonstrated that TCP4 promotes hypocotyl elongation by directly activating the auxin biosynthesis gene *YUC5* (Challa et al., 2016). Together, these findings suggest that GATA25 may promote petiole and hypocotyl elongation through a *TCP4*-dependent growth pathway. Interestingly, despite their elongated petioles, both *GATA25*OX lines exhibited reduced leaf blade areas in leaves 5 and 6 (Fig. 1F). This contrasting effect on petiole elongation and leaf blade expansion is consistent with the established role of *TCP4* in restricting leaf growth by promoting the transition from cell proliferation to differentiation (Palatnik et al., 2003; Schommer et al., 2014; Challa et al., 2019). Furthermore, increased TCP4 activity has been shown to reduce leaf size (Sarvepalli and Nath, 2011). Thus, the simultaneous promotion of petiole elongation and reduction in leaf blade area observed in the *GATA25*OX lines may reflect the distinct roles of *TCP4* in regulating cell elongation and proliferation during leaf development. In contrast to the overexpression phenotypes, the *gata25* mutant exhibited no significant difference in *TCP4* transcript levels or petiole length compared with WT plants (Fig. 1A, D). This lack of an observable phenotype may be explained by functional redundancy among members of the ZIM subfamily, including ZIM-LIKE1 (ZML1) and ZIM-LIKE2 (ZML2) (Shikata et al., 2004). Such redundancy could compensate for the loss of *GATA25* function and maintain *TCP4* expression and petiole growth under the conditions examined.

Our findings further suggest that *GATA25*-mediated transcriptional activation of *TCP4* is influenced by light conditions and photoperiod. GATA25 activated the *TCP4* promoter more strongly under continuous light than under continuous dark conditions in *Arabidopsis* mesophyll protoplasts (Fig. 3B, C; Fig. 4B-E; Supplementary Fig. S7), and under LD conditions than under SD conditions in *N. benthamiana* leaves (Fig. 3G, H; Supplementary Fig. S6). Consistent with these transcriptional responses, *GATA25* induction resulted in a greater relative increase in hypocotyl length under LD conditions than under SD conditions (Fig. 2). These findings are in agreement with a previous study demonstrating that *TCP4* induction promotes hypocotyl elongation in light-grown seedlings but not in dark-grown seedlings (Challa et al., 2016), supporting a role for the GATA25-TCP4 regulatory pathway in light-dependent cell elongation. Interestingly, despite the greater relative contribution of GATA25 to hypocotyl elongation under LD conditions, both mock- and DEX-treated seedlings exhibited longer hypocotyls under SD conditions than under LD conditions (Fig. 2D, F). This difference suggests that the overall hypocotyl growth response to photoperiod is not determined solely by GATA25 activity. Rather, under LD conditions may provide an extended window for *GATA25*-mediated *TCP4* transcriptional activation, enhancing its relative contribution to hypocotyl elongation. Together, these findings suggest that the GATA25-TCP4 regulatory pathway may contribute to the coordination of cell elongation with photoperiod by modulating *TCP4* transcriptional activation in response to light availability.

Our findings reveal that GATA25 binds to a region of the *TCP4* promoter lacking the canonical GATA-binding motif, suggesting that GATA25 may recognize non-canonical DNA sequences. GATA transcription factors typically bind to the conserved (A/T)GATA(A/G) motif through their zinc finger domains (Omichinski et al., 1993; Teakle et al., 2002; Reyes et al., 2004). However, the +61 to +98 region of the *TCP4* promoter, which was identified as a GATA25-binding region by ChIP-qPCR and EMSA (Fig. 4G, H; Fig. S5), does not contain this canonical motif. Instead, this region contains 5’-NATC-3’ and 5’-ATCA(A/G)-3’ sequences, corresponding to 5’-GATN-3’ and 5’-(C/T)TGAT-3’ motifs on the antisense strand (Supplementary Fig. S8). The recognition of non-canonical DNA sequences has been reported for other zinc finger proteins, including White Collar-1 in the fungus *Neurospora crassa*, which binds GATN repeats (Froehlich et al., 2002), and Ash1p in yeast, which recognizes the YTGAT [Y, (C/T)] sequence (Maxon and Herskowitz 2001). Furthermore, previous studies have demonstrated that a single amino acid substitution in the zinc finger loop can alter the DNA-binding specificity of GATA factors (Ravagnani et al., 1997; Starich et al., 1998). Notably, the GATA25 transcription factor possesses a C-X_2_-C-X_20_-C-X_2_-C zinc finger motif (Supplementary Fig. S1D), in contrast to the C-X_2_-C-X_18_-C-X_2_-C motif commonly found in plant GATA transcription factors (Reyes et al., 2004). This structural difference raises the possibility that the extended zinc finger loop of GATA25 contributes to its ability to recognize non-canonical DNA sequences. Together, these findings suggest that GATA25 may exhibit distinct DNA-binding specificity, potentially enabling direct transcriptional regulation of *TCP4* through non-canonical promoter elements.

Our findings suggest that *GATA25*-mediated transcriptional activation of *TCP4* is restricted during the dark period, potentially through post-transcriptional or post-translational regulation of the GATA25 transcription factor. Although *GATA25* transcript levels remained elevated in the GATA25OX lines at ZT16 and ZT20, *TCP4* transcript levels were comparable to those in WT plants (Fig. 5C-G). This contrasts with the increased *TCP4* transcript levels observed at ZT12 (Fig. 1A). Because *GATA25* expression in these lines was driven by the constitutive CaMV 2×35S promoter, these results suggest that elevated *GATA25* transcript abundance alone is insufficient to maintain *TCP4* activation during the dark period. One potential regulator of this light-dependent response is the E3 ubiquitin ligase COP1, which accumulates in the nucleus in darkness and promotes the proteasomal degradation of positive regulators of photomorphogenesis, including HY5 (Osterlund et al., 2000; Holm et al., 2002; Hoecker, 2017). In contrast, light reduces nuclear COP1 accumulation, and photoactivated phyB inhibits COP1/SPA complex activity (von Arnim and Deng, 1994; Pacín et al., 2014; Sheerin et al., 2015). Consistent with a possible role for COP1 in regulating GATA25, our Y2H and BiFC assays demonstrated that GATA25 physically interacts with COP1, with the interaction detected in the nucleus of *N. benthamiana* cells (Fig. 5A, B). Furthermore, GATA25 contains a CCT domain (Supplementary Fig. S1D), and previous studies have shown that the C-terminal region of CO, which contains a CCT domain, is required for COP1 interaction and CO degradation during the night (Jang et al., 2008; Liu et al., 2008; Sarid-Krebs et al., 2015). These observations raise the possibility that COP1 may also regulate GATA25 through a similar mechanism.

Based on these findings, we propose a model in which COP1 may negatively regulate GATA25 protein stability during the dark period, limiting *GATA25*-mediated transcriptional activation of *TCP4* (Fig. 6). This proposed mechanism is consistent with the reported daily expression patterns of *GATA25*, which peaks around subjective dawn (Manfield et al., 2007), and *TCP4*, which peaks between ZT10 and ZT13 under LD conditions (Kubota et al., 2017). Thus, COP1-mediated regulation of GATA25 may contribute to the coordination of *TCP4* expression with the daily light/dark cycle.

**Fig. 6.**
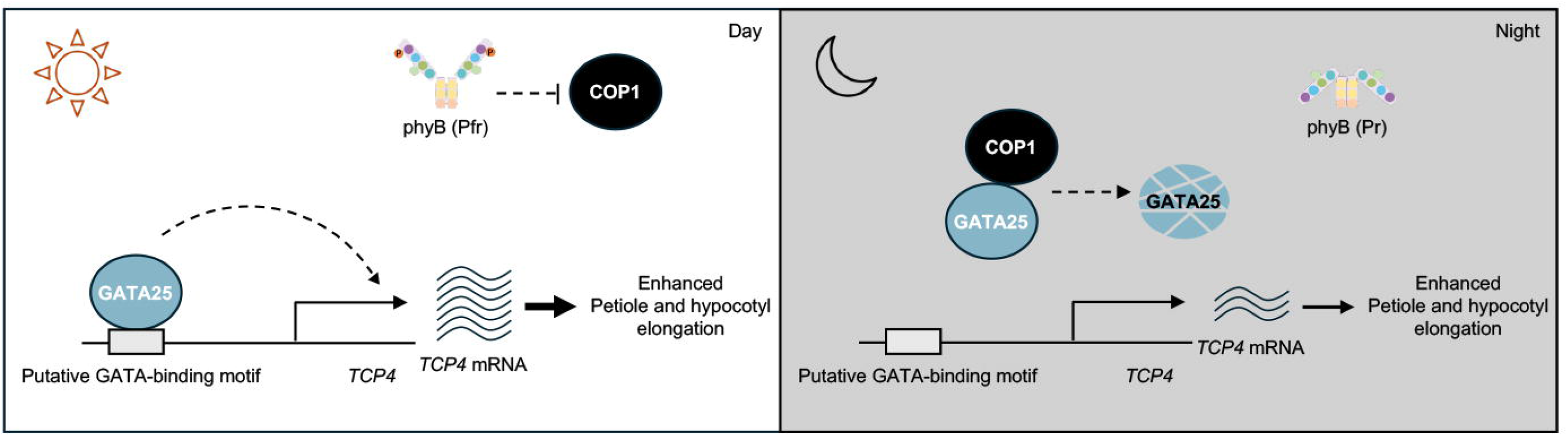
Proposed model for the light/dark-dependent regulation of *TCP4* by GATA25 transcription factor. During the day, photoactivated phytochrome B (phyB, Pfr form) suppresses COP1 activity, allowing GATA25 transcription factor to bind the putative GATA-binding motif of the *TCP4* promoter and activate *TCP4* transcription, which promotes petiole and hypocotyl elongation. During the night, COP1 is activated and interacts with GATA25 transcription factor, and GATA25 transcription factor is proposed to be destabilized, which reduces the activation of *TCP4* transcription. Dashed arrows indicate proposed regulatory steps that were not directly tested in this study.

## Author contributions

KK and WCK: designed and coordinated the research; KK and JS: conducted most of the experiments and data analysis; KK: wrote the first version of the manuscript, and other authors contributed later to improve it and approved the final version for submission. All authors contributed to the article and approved the submitted version.

## Data availability

All data needed to evaluate the conclusions in the paper are present in the paper and its supplementary data.

## Supplementary data

**Fig. S1.** Constructs and generation of the *GATA25*-overexpressing transgenic lines and *gata25* mutant. (A) Schematic diagram of the *GATA25*/pTK-BMLC binary vector. Scale bar, 300 bp. (B) Schematic diagram of the *GATA25*/pKSE401 binary vector and the gRNA sequence. gRNA scaffold, guide RNA scaffold sequence; *Kan^R^*, kanamycin resistance gene; P_U6-26_, U6-26 promoter; *zCas9*, *Zea mays* codon-optimized *Cas9* sequence. Scale bar, 1000 bp. (C) Position of the gRNA target and protospacer adjacent motif (PAM) in the *GATA25* gene. Scale bar, 150 bp. (D) Domain structure of GATA25 transcription factor showing the TIFY domain, the CCT domain, and the GATA-type zinc finger with C-X_2_-C-X_20_-C-X_2_-C motif. Scale bar, 75 amino acids. (E) Nucleotide and deduced amino acid sequences of the target region in the *gata25* mutant, showing a 1 bp insertion that causes a frameshift and a premature stop codon. Scale bar, 150 bp. (F) Predicted truncated protein in the *gata25* mutant Scale bar, 75 amino acids.

**Fig. S2.** Selection of the *GATA25*-overexpressing transgenic lines. (A) Representative photographs of 4-week-old WT, *gata25*, and independent *GATA25*-overexpressing transgenic lines. Scale bar, 7 cm. (B) Semi-quantitative RT-PCR analysis of *GATA25* transcripts in the transgenic lines. *ACT8* was used as the internal control. (C) Quantification of band intensity (*GATA25*/*ACT8*) using ImageJ software.

**Fig. S3.** PCR-based validation of the transgenic lines and the *gata25* mutant. (A) Schematic diagram of the *GATA25*/pTK-BMLC binary vector. The positions of the 35S minimal promoter_F and *GATA25*_mid_R primers used for detection of the *GATA25* expression cassette are indicated. Scale bar, 300 bp. (B) Schematic diagram of the *GATA25*/pKSE401 binary vector. The *GATA25*-targeting gRNA and the positions of the pKSE401_F and pKSE401_R primers used for detection of the *zCas9*-containing cassette are indicated. Scale bar, 300 bp. (C) PCR-based validation of *GATA25*-overexpressing lines (*GATA25*OX #3 and #13) and the *gata25* mutant. *ACT8* gene was amplified as a genomic DNA control using *ACT8*_F/R primers, yielding a 403 bp PCR product. The *GATA25* overexpression cassette was detected using the 35S minimal promoter_F and *GATA25*_mid_R primers, yielding a 1,043 bp PCR product. The pKSE401-derived cassette was detected using pKSE401_F/R primers, yielding a 630 bp PCR product. The absence of this product in the *gata25* mutant indicates that the mutant no longer carries the *zCas9* cassette. M, DNA size marker.

**Fig. S4.** Generation of 2×35S::*GATA25*-mRFP transgenic lines. (A) Schematic diagram of the *GATA25*/pTK-mRFP-BMLC binary vector. The *GATA25* CDS without the stop codon was placed under the control of the CaMV 2×35S promoter and fused in-frame to mRFP. Scale bar, 300 bp. (B) Representative photographs of 5-day-old seedlings of WT, 2×35S::*GATA25*-mRFP #1, and 2×35S::*GATA25*-mRFP #4. Scale bar, 2 mm. (C) *GATA25* transcript levels in WT, 2×35S::*GATA25*-mRFP #1, and 2×35S::*GATA25*-mRFP #4. Transcript levels were normalized to *PP2A*. Data are presented as mean ± SD of three biological replicates. Different letters indicate significant differences (one-way ANOVA with Tukey’s HSD test, *p* < 0.05).

**Fig. S5.** Identification of GATA25-binding regions in the *TCP4* promoter by EMSA. (A) SDS-PAGE analysis of the purified His_6_-MBP and His_6_-GATA25 recombinant protein. M, protein size marker; S, soluble protein; P, Ni^2+-^NTA-based purified protein. (B) Schematic diagram of the *TCP4* promoter fragment used for EMSA. Scale bar, 150 bp. (C) EMSA of the *TCP*4-1, *TCP*4-2, and *TCP*4-3 fragment with increasing amounts of His_6_-GATA25 recombinant protein separated on a 0.6% agarose gel. (D) Schematic diagram of the *TCP4*-3-1-1 and *TCP4*-3-1-2 fragment. Scale bar, 150 bp. (E, F) EMSA of the biotin-labeled *TCP4*-3-1-1 (E) and *TCP4*-3-1-2 (F) probes with increasing amounts of His_6_-GATA25 recombinant protein. His_6_-MBP recombinant protein was used as the negative control. Increasing numbers of “+” symbols indicate increasing protein concentrations, with “++++” corresponding to 200 nM.

**Fig. S6.** GUS activity in agroinfiltrated *N. benthamiana* leaves under different photoperiod conditions. The effector and reporter constructs shown in Figure 3D were co-infiltrated into *N. benthamiana* leaves to compare GUS activity under SD and LD conditions. GUS activity was expressed in picomoles of 4-methylumbelliferone produced per minute per milligram of protein (pmol 4-MU/min/mg protein). Data are presented as mean ± SD of six biological replicates. *p* values were determined by Welch’s *t*-test (\**p* < 0.05, \*\**p* < 0.005, and \*\*\**p* < 0.001).

**Fig. S7.** Relative GUS activity in *Arabidopsis* mesophyll protoplasts under light or dark conditions. The effector and reporter constructs shown in Figure 4A were co-transfected into *Arabidopsis* mesophyll protoplasts to compare GUS activity under continuous dark and continuous light conditions following treatment with 10 μM DEX. Data are presented as mean ± SD of three biological replicates. *p* values were determined by Welch’s *t*-test (\**p* < 0.05, \*\**p* < 0.005, and \*\*\**p* < 0.001).

**Fig. S8.** Sequence analysis of *TCP4* promoter fragments used for EMSA-based mapping of GATA25-responsive regions. The *TCP4* promoter was divided into *TCP4*-1, *TCP4*-2, and *TCP4*-3 fragments for initial screening. *TCP4*-3 was subsequently subdivided into *TCP4*-3-1, *TCP4*-3-2, and *TCP4*-3-3, and *TCP4*-3-1 was further divided into *TCP4*-3-1-1 and *TCP4*-3-1-2 to narrow down the GATA25-responsive region. Putative GATA-binding motifs are color-coded within each fragment. The analyzed motifs comprised 5’- (A/T)GATA(G/A)-3’, 5’-CGATA(G/A)-3’, 5’-TGATAG-3’, 5’-(A/C)GATAG-3’, 5’-(C/T)TGAT-3’, and 5’-GATN-3’, as well as the corresponding reverse-complement sequences.

Table S1. List of primers used for vector construction and validation of DNA presence in this study

Table S2. List of primers used for gene expression analysis and ChIP-qPCR enrichment assays in this study

Table S3. List of primers used for amplification of DNA fragments for EMSA in this study

